# Cefiderocol heteroresistance associated with mutations in TonB-dependent receptor genes in *Pseudomonas aeruginosa* of clinical origin

**DOI:** 10.1101/2024.01.30.578008

**Authors:** Stephanie L. Egge, Samie A. Rizvi, Shelby R. Simar, Manuel Alcalde, Jose R.W. Martinez, Blake M. Hanson, An Q. Dinh, Rodrigo P. Baptista, Truc T. Tran, Samuel A. Shelburne, Jose M. Munita, Cesar A. Arias, Morgan Hakki, William R. Miller

**Author notes:** <u>Corresponding Author:</u> William Miller, Division of Infectious Diseases, Houston Methodist Hospital, 6560 Fannin St, Scurlock Tower Suite 1540, Houston, TX 77030. Current Address/Affiliation: Department of Medicine, Division of Infectious Diseases, Oregon Health and Science University, 3181 SW Sam Jackson Park Road, Portland, OR 97239.

## Abstract

The siderophore-cephalosporin cefiderocol(FDC) presents a promising treatment option for carbapenem-resistant (CR) *P. aeruginosa* (PA). FDC circumvents traditional porin and efflux mediated resistance by utilizing TonB-dependent receptors (TBDRs) to access the periplasmic space. Emerging FDC resistance has been associated with loss of function mutations within TBDR genes or the regulatory genes controlling TBDR expression. Further, difficulties with antimicrobial susceptibility testing (AST) and unexpected negative clinical treatment outcomes have prompted concerns for heteroresistance, where a single lineage isolate contains resistant subpopulations not detectable by standard AST. This study aimed to evaluate the prevalence of TBDR mutations among clinical isolates of *P. aeruginosa* and the phenotypic effect on FDC susceptibility and heteroresistance. We evaluated the sequence of *pirR*, *pirS*, *pirA*, *piuA* or *piuD* from 498 unique isolates collected before the introduction of FDC from 4 clinical sites in Portland, OR (1), Houston, TX (2), and Santiago, Chile (1). At some clinical sites, TBDR mutations were seen in up to 25% of isolates, and insertion, deletion, or frameshift mutations were predicted to impair protein function were seen in 3% of all isolates (n=15). Using population analysis profile testing, we found that *P. aeruginosa* with major TBDR mutations were enriched for a heteroresistant phenotype and undergo a shift in the susceptibility distribution of the population as compared to susceptible strains with wild type TBDR genes. Our results indicate that mutations in TBDR genes predate the clinical introduction of FDC, and these mutations may predispose to the emergence of FDC resistance.

## Introduction

Multi-drug resistant (MDR) *Pseudomonas aeruginosa* (PA) is a priority antimicrobial resistant (AMR) pathogen with an estimated mortality impact of 2,700 hospitalized US patients per year (1, 2). Moreover, rates of MDR *P. aeruginosa* in hospitalized and critically ill patients are rising, increasing dependence on last line agents such as the novel cephalosporin cefiderocol (FDC). Emerging clinical data have raised concerns about the reliability of FDC antimicrobial susceptibility testing (AST) and the translation of *in vitro* susceptibility data to clinical efficacy (3, 4). Despite high rates of laboratory-reported susceptibility in carbapenem-resistant pathogens, a numerical increase in mortality was seen among patients randomized to cefiderocol based therapy versus polymyxin based combination therapy in a phase 3 study (5). Further, clinical isolates of *P. aeruginosa* that test susceptible by routine AST have been described to rapidly develop cefiderocol resistance upon drug exposure (6–8). These data emphasize a need to define microbiologic and clinical risk factors for cefiderocol treatment failure.

Bacterial heteroresistance has been suggested as one potential explanation for the discordance in microbiologic susceptibility data and clinical outcomes (9). Heteroresistance describes a phenomenon whereby minority subpopulations of an isolate display an antibiotic-resistant phenotype below the detectable limits of standard clinical microbiology techniques. Under antimicrobial pressure, these low frequency resistant subpopulations survive antibiotic killing and continue to expand, potentially leading to recurrent infection and the emergence of resistance and associated treatment failure (10, 11). Heteroresistance to cefiderocol has been reported in association with *P. aeruginosa* infections where an initial isolate tested susceptible by standard antimicrobial susceptibility testing, but cefiderocol non-susceptibility later emerged (12). However, the mechanisms underlying this phenotype remain poorly understood.

As a catechol conjugated cephalosporin, cefiderocol functions under the guise of an iron-siderophore and circumvents traditional drug resistance pathways by binding iron and utilizing TonB-dependent receptors (TBDR) to enter the periplasmic space (13–16). In *P. aeruginosa*, there are two putative TBDRs that contribute to the vast majority of cefiderocol uptake, namely PirA and PiuA (with some strains possessing the ortholog PiuD instead of PiuA) (15, 16). Import of catechol siderophores by these TBDRs leads to activation of the PirRS two-component sensor system that promotes a positive-feedback loop on transcription of *pirA* and *piuA/piuD* resulting in increased production of the catechol-siderophore receptors (17). Expression of *pirA* is tightly controlled via phosphorylation of the response regulator PirR via the histidine kinase PirS. Of note, there is a basal level of constitutive expression of *piuA* even in the absence of PirS.

Clinical and *in vitro* data suggest resistance to catechol-conjugated antibiotics emerges when both PirA and PiuA/D become non-functional either via direct mutations in *piuA/D* and *pirA* or via mutations in *pirS* or *pirR* that decrease the expression of these TBDRs (6, 8, 15). However, while *in vitro* minimum inhibitory concentrations required for *P. aeruginosa* killing are elevated with the loss of a single system (e.g. PirA or PiuA), overt resistance to catechol conjugated antibiotics does not emerge when single transporter isogenic *P. aeruginosa* knockouts have been generated (18). Thus, underlying mutations in these pathways may not be recognized using conventional AST despite their serving as a potential first step in the emergence of cefiderocol resistance.

To date, no study has investigated the clinical prevalence of mutations in genes encoding the PirA, PiuA/D and PirRS proteins among clinical isolates of *P. aeruginosa*. Here, we investigate the prevalence of mutations in these genes across geographically diverse sets of clinical isolates and assess the impact of these mutations on cefiderocol susceptibility and their association with heteroresistance.

## Methods

### Isolate source and identification of polymorphisms in TBDR genes

We analyzed whole genome sequence data for clinical isolates of *P. aeruginosa* previously collected from 2006-2018 from four institutions: an urban hospital in Houston, TX (n=212), a referral cancer hospital in Houston, TX (n=107), an urban hospital in Portland, OR (n=80), and an urban hospital in Santiago, Chile (n=99) (19, 20). Isolates were recovered from variable clinical sources (e.g. sputum, urine, blood), and patients had no prior exposure to FDC (as FDC was not clinically available at the time of PA isolate collection). The isolates collected from the general hospitals in Houston, TX and Santiago, Chile were all carbapenem-resistant, while those from the Portland, OR general hospital and the Houston, TX cancer hospital were a mix of carbapenem resistant and susceptible isolates. For all four libraries, whole genome sequence data was used to evaluate for mutations of genes encoding TBDR component proteins previously implicated in decreased susceptibility to FDC: *pirA*, *pirR*, *pirS*, and *piuA* or *piuD*. Sequences were assessed for the prevalence of: 1) insertion, deletion, frameshifts, 2) early stop codon, and 3) single nucleotide polymorphisms. Genomes were assembled with SPAdes (version 3.15.2), annotated with RAST, and then differences in both nucleotide and predicted amino acid sequence changes were determined by pairwise alignment using *P. aeruginosa* PAO1 as a reference.

### Antimicrobial susceptibility testing

Phenotypic studies were performed from the Houston, TX general hospital isolate library as these isolates were previously banked and available for further laboratory testing. From this library, all isolates found to have probable loss of function mutations in the TBDR pathway proteins (e.g. frameshift, insertion, deletion, early stop) were obtained for microbiologic resistance analyses. *P. aeruginosa* PAO1 and carbapenem-resistant clinical isolates from the high-risk ST111 and ST235 lineages with wild type TBDR gene sequences were used as controls. Each isolate was grown on fresh cetrimide agar (Oxoid) from purified stocks stored in Brucella broth with 15% glycerol at −80° C. Cefiderocol susceptibility testing was performed using broth microdilution in iron-depleted Mueller-Hinton (ID-MH, BBL Mueller Hinton II cation-adjusted, Becton Dickinson) media prepared in accordance with the Clinical and

Laboratory Standards Institute M100 Appendix I (using Chelex 100 resin, BioRad Laboratories) and verified to have a final iron concentration ≤ 0.03 mg/L (MilliporeSigma MQuant Iron Test Kit)(21, 22). Broth microdilution plates were made using ID-MH to create a gradient of FDC concentrations 0.5 to 32 ug/mL. A 0.5 McFarland standard was measured, diluted 1:100 in ID-MH, and 50 µL was inoculated into each well to achieve a final inoculum of 5×10^5 CFU/mL. Plates were incubated for 20 hours after which minimum inhibitory concentrations(MICs) were recorded. Kirby-Bauer disk diffusion testing was performed on standard Mueller-Hinton agar (BBL Mueller Hinton II agar, Becton Dickinson). For each isolate, a 0.5 McFarland standard was plated and a single FDC drug eluting disc (HardyDisks™ AST cefiderocol 30 µg) was placed at the center of the cultured plate. Plates were incubated for 48 hours at 37° C with disk diameter measurements taken at 24-hour and 48-hour time points.

### Population analysis profile (PAP) and calculation of area under the curve (AUC) measurements

Initial screening was performed using a single colony selected at random and grown overnight in ID-MH. For each isolate, plates were prepared with MH agar supplemented to attain a gradient of FDC concentrations: 0, 4, 8, and 16 µg/mL (1x, 2x, and 4x the cutoff for susceptible). Overnight cultures were diluted in phosphate buffer saline (PBS) to create a series of ten-fold dilutions and plated onto a labeled grid using 50 µL aliquots in technical triplicate per plate. Plates were incubated and read at 24 and 48 hours. Colony forming units (CFUs) per mL were counted per each corresponding section and averaged across technical triplicates. CFU values were log-transformed and normalized to growth relative to each isolate’s corresponding 0 µg/mL FDC plate (Equation 1) to obtain Δlog values to calculate relative survival under different FDC concentration exposures.

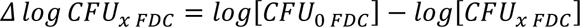

where *x* corresponds to CFU values obtained at either 4, 8, 16 µg/mL FDC exposure

**Equation 1**. PAP Isolate Growth Normalization

Using previously published criteria, susceptible was defined as a ≥ 6-log_10_ drop in CFU/mL at 4 µg/mL, heteroresistance was defined as less than 50% isolate survival at 4 µg/mL with greater than 0.0001% survival at 8 µg/mL (when compared to 0 µg/mL FDC growth), and resistance was defined as > 50% isolate survival at 8 µg/mL or higher (9, 23).

After initial screening, PAP-AUC was performed in triplicate using a controlled inoculum of 0.5 McFarland (approximately 2×10^8^ CFU/mL) for all four TBDR mutants identified catalogued with a heteroresistant phenotype, as well as for 3 TBDR mutants with a susceptible phenotype and 2 susceptible non-mutant controls. Using the Δlog values, area under the curve (AUC) was calculated with GraphPad Prism with a baseline of −10 using a trapezoidal method between any two subsequent points with inclusion of all Y increases that exceeded 0. A baseline of −10 was chosen as the maximum log change observed in our study was −9.15. Welch’s t-test was run to assess differences in initial inocula and AUCs. Differences in colony counts across the different antibiotic concentrations in the PAP experiments were analyzed using ANOVA with Dunnett’s test for multiple comparisons and PAO1 as reference. Linear regression was used to compare trends between KB and PAP-AUC results.

### Validation of agar dilution PAP using ID-MH broth

Isolates were grown in ID-MH overnight and adjusted to a controlled inoculum of 0.5 McFarland. The CFU of the controlled inoculum was verified by plating a serial dilution of the initial McFarland to an antibiotic free plate for pre-incubation colony counts. This standard was then diluted 1:10 and approximately 1×10^7^ CFU/mL was inoculated into ID-MH broth with FDC concentrations at 0, 4, 8, and 16 µg/mL and grown at 37° C in a rotary incubator for 24 hours. At 24 hours, broth cultures were centrifuged and washed twice in PBS to remove the broth-antibiotic supernatant. Pellets were then resuspended in PBS and serial ten-fold dilutions were plated on antibiotic free Mueller-Hinton agar. Plates were incubated at 37° C for 24 hours after which colonies could be counted to quantify survival, and AUC was determined as above. The heteroresistant isolates C1814 and C4593 were tested to represent the highest and lowest AUC by agar PAP testing, and the susceptible non-TBDR mutant C1273 and laboratory strain PAO1 served as controls. All assays were performed in technical and biological triplicates.

### Evaluation of population MIC distribution

All four heteroresistant isolates were tested, and the susceptible non-TBDR mutant C1273 and laboratory strain PAO1 served as controls. Isolates were grown in ID-MH overnight and adjusted to a controlled inoculum of 0.5 McFarland in fresh ID-MH before exposure to cefiderocol at 0, 4, and 16 µg/mL. After growth at 37° C in a rotary incubator for 24 hours, broth cultures were centrifuged and washed twice in PBS to remove the broth-antibiotic supernatant. Pellets were then resuspended in PBS and plated on antibiotic free MH agar. Three unique colonies (growth permitting) were collected at random from each plate corresponding to the different antibiotic concentrations and incubated on MH agar for 24 hours at 37°C. These colonies were subcultured one additional time in the absence of antibiotic, and then FDC MICs were determined using broth microdilution in triplicate for each subculture isolate. Overnight growth in antibiotic was repeated in biological triplicate. GraphPad Prism was used to assess statistical differences in population MIC distributions for each isolate using two-way ANOVA with Dunnett’s test to correct for multiple comparisons.

## Results

### Prevalence of mutations in the PirRS and PiuA/D pathways from clinical isolates of *P. aeruginosa*

To evaluate the frequency of mutations in the PirRS and PiuA/D pathways, we utilized a collection of isolates for which sequencing data was available from four distinct geographic locations and patient populations: (1) a general hospital in Houston, Texas^19^ (2) a cancer hospital in Houston, Texas, (3) a general hospital in Portland, Oregon^20^, and (4) an urban hospital in Santiago, Chile (19, 20). All isolates were collected prior to the introduction of cefiderocol. Overall, single nucleotide polymorphisms (SNPs) were present in between 5% to 30% of the evaluated TBDR components across all isolate libraries (**Figure 1A**). Major *pirRS* and *piuA/D* mutations likely to have a significant impact on protein function, defined as a nucleotide change leading to an amino acid insertion or deletion, frameshift, or premature stop codon, were detected only in hospitals from Houston, TX and not in isolates from Portland, Oregon or Chile. In Houston general hospital isolates, *pirR* had the highest frequency of probable loss of function mutations with 9 isolates (4.7%) exhibiting insertion, deletion, or frameshift mutations, and 1 (0.5%) harboring an early stop codon (**Figure 1B**). For *piuA/piuD*, approximately 1% of isolates had insertion, deletion, or frameshift mutations. Isolates obtained from the Houston cancer hospital had a similar distribution of changes in TBDR genes to those from the general hospital within the same city. Nearly 2% of Houston, TX cancer hospital isolates had insertion, deletion, frame shift, or early stop codon mutations, and one isolate had an early stop mutation within *pirR* (**Figure 1B**).

**Figure 1.**
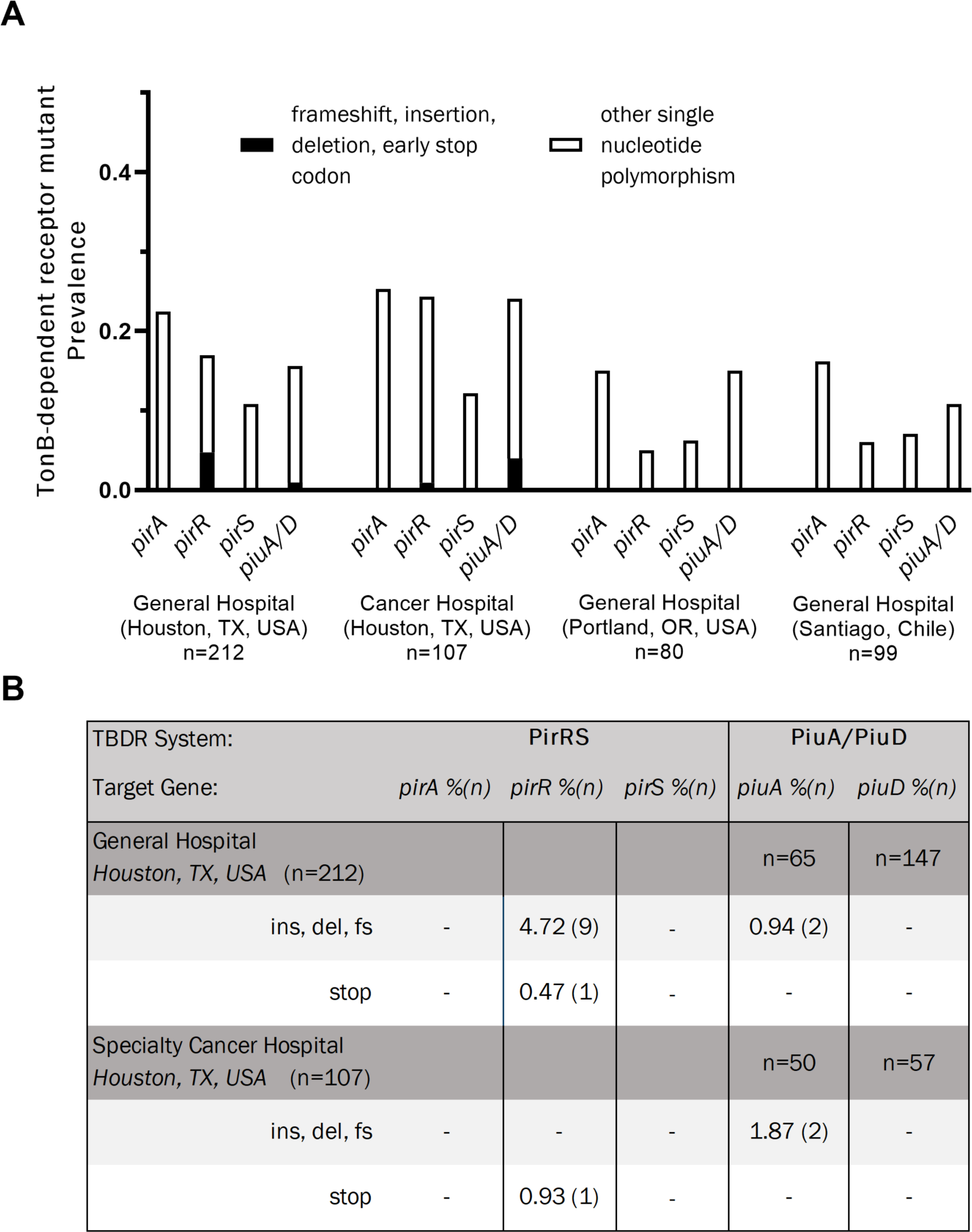
TonB dependent receptor mutant prevalence across four clinical populations of *Pseudomonas aeruginosa.* **A)** Prevalence of mutations in the genes encoding the PirA and PiuA/D cefiderocol import s. **B)** Distribution of major mutations, defined as those resulting in a frameshift (fs), insertion (ins), deletion (del), and/or early stop codon.

### Characterization of major TBDR mutations

To characterize the impact of the identified major mutations in TDBR genes on susceptibility to cefiderocol, all 12 isolates harboring major mutations in the Houston, TX general hospital collection underwent testing by both broth microdilution in ID-MH media and by disk diffusion testing (**Table 1**). The diversity in the observed number of sequence types (STs) suggested that isolates with major TBDR mutations were not representative of clonal expansion, although 5 of the 12 isolates belonged to one of three high-risk clonal lineages (ST235, n=1; ST111, n=2; ST274, n=2). The laboratory strain *P. aeruginosa* PAO1 and five carbapenem-resistant isolates from the high-risk lineages ST235 and ST111 with wild type TBDR sequences were used as controls. Broth microdilution results of major TBDR mutants characterized 1 isolate as resistant (C1408, FDC MIC 16 µg/mL) and 1 as intermediate (C1418, FDC MIC 8 µg/mL) by CLSI susceptibility criteria (**Table 1**). Notably, the resistant isolate possessed a variant of the OXA-15 beta-lactamase with a G150S amino acid substitution, while the intermediate isolate had the PDC-205 variant. Both isolates were also non-susceptible by disk diffusion with zone diameters of 13 mm and 12 mm for C1408 and C1418, respectively. The remaining 10 of 12 TBDR isolates and all control isolates tested susceptible by both broth microdilution and KB disk diffusion (**Table 1**).

**Table 1.**
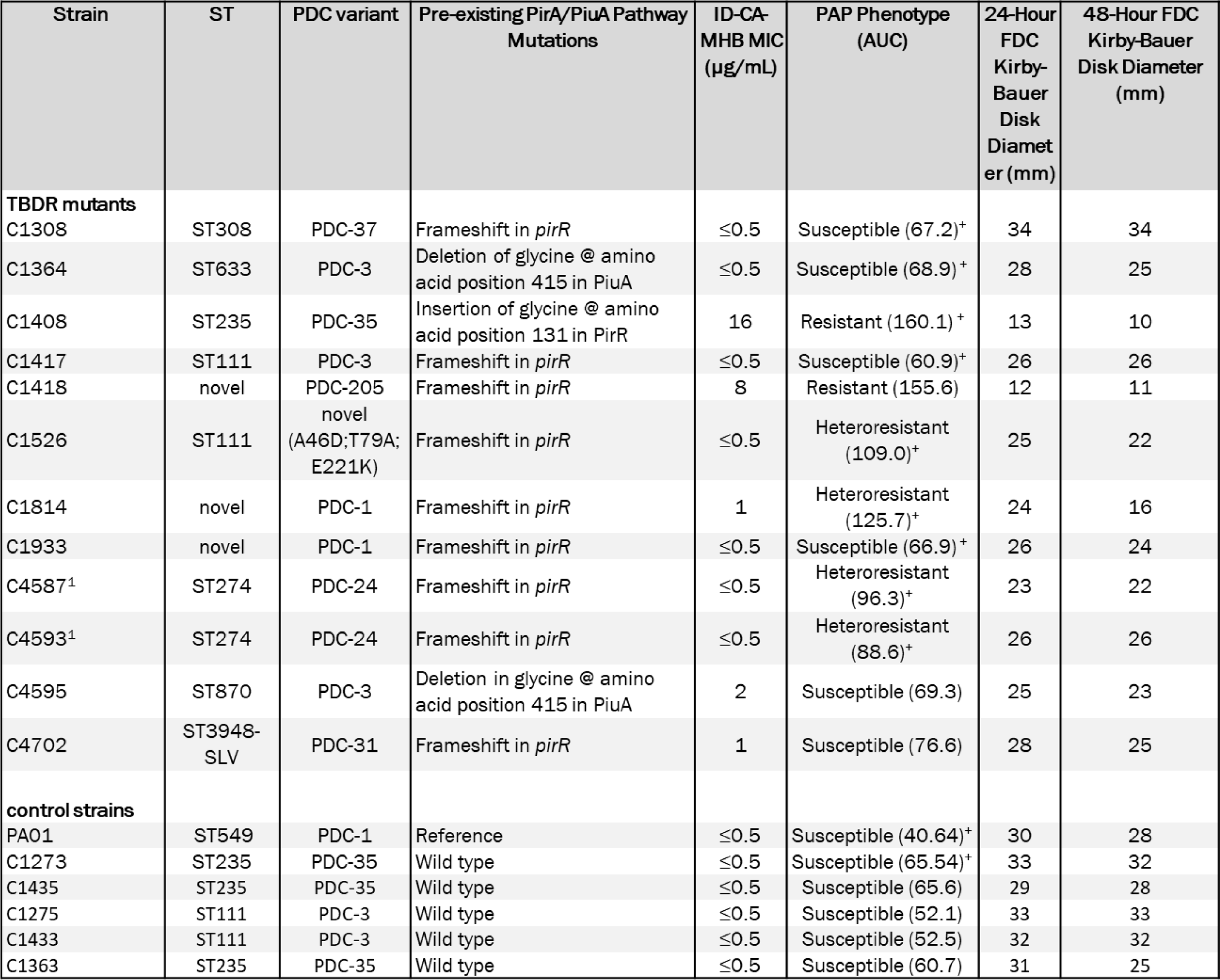
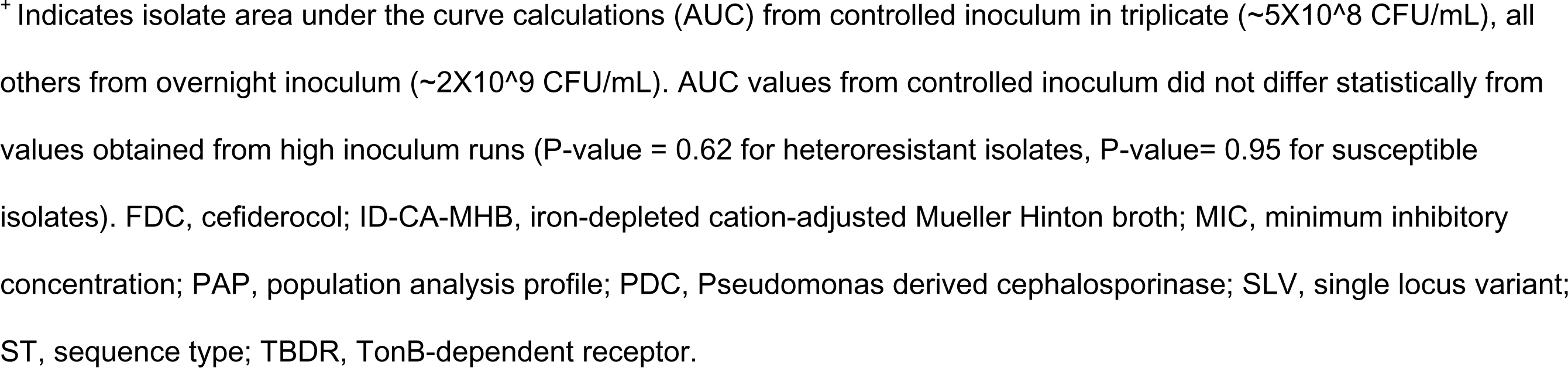
Summary of TBDR mutant backgrounds including specific TBDR mutations, sequence type, additional beta-lactamase properties, and cefiderocol broth microdilution susceptibility test results.

| Strain | ST | PDC variant | Pre-existing PirA/PiuA Pathway Mutations | ID-CA-MHB MIC (µg/mL) | PAP Phenotype (AUC) | 24-Hour FDC Kirby-Bauer Disk Diameter (mm) | 48-Hour FDC Kirby-Bauer Disk Diameter (mm) |
| --- | --- | --- | --- | --- | --- | --- | --- |
| **TBDR mutants** |  |  |  |  |  |  |  |
| C1308 | ST308 | PDC-37 | Frameshift in *pirR* | ≤0.5 | Susceptible (67.2)<sup>+</sup> | 34 | 34 |
| C1364 | ST633 | PDC-3 | Deletion of glycine @ amino acid position 415 in PiuA | ≤0.5 | Susceptible (68.9)<sup>+</sup> | 28 | 25 |
| C1408 | ST235 | PDC-35 | Insertion of glycine @ amino acid position 131 in PirR | 16 | Resistant (160.1)<sup>+</sup> | 13 | 10 |
| C1417 | ST111 | PDC-3 | Frameshift in *pirR* | ≤0.5 | Susceptible (60.9)<sup>+</sup> | 26 | 26 |
| C1418 | novel | PDC-205 | Frameshift in *pirR* | 8 | Resistant (155.6) | 12 | 11 |
| C1526 | ST111 | novel (A46D;T79A;E221K) | Frameshift in *pirR* | ≤0.5 | Heteroresistant (109.0)<sup>+</sup> | 25 | 22 |
| C1814 | novel | PDC-1 | Frameshift in *pirR* | 1 | Heteroresistant (125.7)<sup>+</sup> | 24 | 16 |
| C1933 | novel | PDC-1 | Frameshift in *pirR* | ≤0.5 | Susceptible (66.9)<sup>+</sup> | 26 | 24 |
| C4587<sup>1</sup> | ST274 | PDC-24 | Frameshift in *pirR* | ≤0.5 | Heteroresistant (96.3)<sup>+</sup> | 23 | 22 |
| C4593<sup>1</sup> | ST274 | PDC-24 | Frameshift in *pirR* | ≤0.5 | Heteroresistant (88.6)<sup>+</sup> | 26 | 26 |
| C4595 | ST870 | PDC-3 | Deletion in glycine @ amino acid position 415 in PiuA | 2 | Susceptible (69.3) | 25 | 23 |
| C4702 | ST3948-SLV | PDC-31 | Frameshift in *pirR* | 1 | Susceptible (76.6) | 28 | 25 |
| **control strains** |  |  |  |  |  |  |  |
| PA01 | ST549 | PDC-1 | Reference | ≤0.5 | Susceptible (40.64)<sup>+</sup> | 30 | 28 |
| C1273 | ST235 | PDC-35 | Wild type | ≤0.5 | Susceptible (65.54)<sup>+</sup> | 33 | 32 |
| C1435 | ST235 | PDC-35 | Wild type | ≤0.5 | Susceptible (65.6) | 29 | 28 |
| C1275 | ST111 | PDC-3 | Wild type | ≤0.5 | Susceptible (52.1) | 33 | 33 |
| C1433 | ST111 | PDC-3 | Wild type | ≤0.5 | Susceptible (52.5) | 32 | 32 |
| C1363 | ST235 | PDC-35 | Wild type | ≤0.5 | Susceptible (60.7) | 31 | 25 |

<sup>+</sup> Indicates isolate area under the curve calculations (AUC) from controlled inoculum in triplicate ( $\sim 5 \times 10^8$ CFU/mL), all others from overnight inoculum ( $\sim 2 \times 10^9$ CFU/mL). AUC values from controlled inoculum did not differ statistically from values obtained from high inoculum runs (P-value = 0.62 for heteroresistant isolates, P-value= 0.95 for susceptible isolates). FDC, cefiderocol; ID-CA-MHB, iron-depleted cation-adjusted Mueller Hinton broth; MIC, minimum inhibitory concentration; PAP, population analysis profile; PDC, Pseudomonas derived cephalosporinase; SLV, single locus variant; ST, sequence type; TBDR, TonB-dependent receptor.

### Evaluation of heteroresistance by population analysis profiling

Heteroresistance, or the presence of a subpopulation of bacterial cells with a reduced antimicrobial susceptibility, has been described for β-lactam antibiotics including cefiderocol, although the specific mechanisms responsible for this phenotype have not been identified (24). We sought to evaluate if the presence of major mutations in the TBDRs mediating cefiderocol uptake were associated with heteroresistance using an agar dilution population analysis profile (PAP) method. Both isolates identified as non-susceptible by standard susceptibility criteria were also identified as resistant using PAP. Among the remaining 10 TBDR mutants, PAP studies identified 4 isolates as heteroresistant and 6 isolates as susceptible. All 5 susceptible controls tested susceptible per agar dilution PAP tests (**Supplemental Figure 1**), and there were no significant differences in the starting inoculums between the susceptible and heteroresistant strains (3.62 X 10^9^ CFU/mL versus 2.21 X 10^9^ CFU/mL, p=0.30).

Area under the curve (AUC) analyses showed a statistically significant difference in the AUC of susceptible, resistant, and heteroresistant isolates, and these results were consistent regardless of inoculum. The mean AUC for heteroresistant and for susceptible isolates was 111.2 (95% CI 89.29-133.2) and 63.5 (CI 95% 49.88-75.19), respectively. This mean difference was statistically significant (two-sided t-test, p=0.0087). The PAP-AUC results were similar when performed with a controlled inoculum of approximately 2×10^8^ CFU/mL on a subset of the isolates. Controlled inoculum studies performed in triplicate showed a mean AUC of 104.9 (CI 95% 85.72-119.4) for the heteroresistant isolates, and of 61.7 (CI 95% 48.68-71. 07) for the susceptible isolates (p=0.0061, **Supplemental Figure 2**).

### Heteroresistance is not an artifact of agar iron content

Previous studies have suggested that the iron in the MH agar media is bound in such a way that the locally available concentrations are low and mimic an iron-deficient environment (25, 26). However, there are potential concerns that the presence of excess iron may lead to difficulties in interpreting cefiderocol susceptibility (3, 27, 28). To verify that the differences in population survival seen on agar dilution PAP were not an artifact of the iron content of the media, we performed killing assays across the same cefiderocol concentrations using a controlled inoculum in ID-MH broth. Two heteroresistant isolates (C1814 and C4593) were chosen for testing as these isolates represented those with the highest and lowest AUCs in the heteroresistant group, as well as the laboratory strain PAO1 and a susceptible clinical control with wild type TBDR genes (C1273).

Both C1814 and C4593 maintained a heteroresistant phenotype, with a 3.22 and 5.28 log_10_ reduction in CFU/mL at 16 µg/mL, respectively (**Figure 2**). Both the clinical control C1273 and laboratory strain PAO1 remained fully susceptible, with a greater than 6 log_10_ decrease in CFU/mL at 16 µg/mL. Compared to PAO1, isolate C1814 had a statistically significant increase in growth at 8 µg/mL (p<0.0001) and 16 µg/mL (p<0.0001). While the mean CFU/mL of isolate C4593 remained higher than the 6 log_10_ cutoff for heteroresistance, the growth difference as compared to both control strains did not reach statistical significance (**Figure 2B, Supplemental Figure 3**).

**Figure 2.**
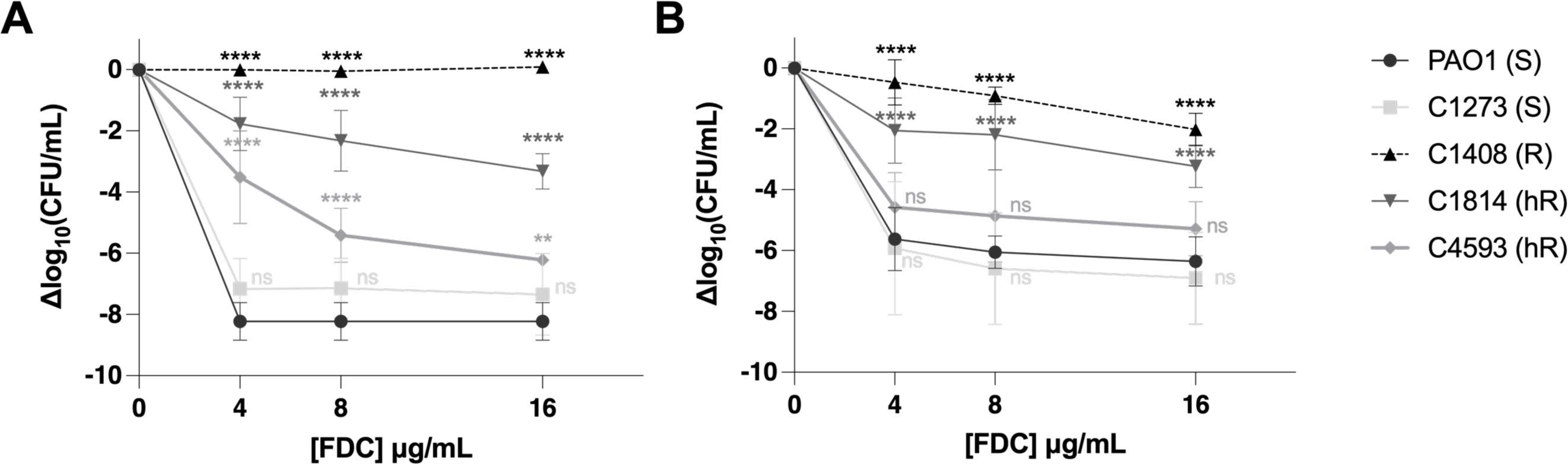
Heteroresistance phenotype associated with TBDR mutations is present independent of the iron state of growth media upon cefiderocol exposure. **A)** Population analysis profile performed by agar dilution on ascending concentrations of cefiderocol in Muller-Hinton agar. **B)** Population analysis performed by growth of strains with ascending cefiderocol concentrations in iron-depleted Mueller-Hinton broth. Error bars represent standard deviation of three independent runs. Asterix above each point represents the statistical difference from PAO1 at each indicated concentration. **, p<0.01; ****, p<0.0001; ns, non-significant.

### Heteroresistance is associated with a change in the population MIC distribution after exposure to cefiderocol

To assess whether the heteroresistance phenotype was associated with changes in susceptibility to cefiderocol, we sampled isolates from the original strain populations grown in ID-MH broth before and after exposure to cefiderocol at 4 µg/mL (susceptible breakpoint) and 16 µg/mL (resistant breakpoint). Growth of subcultured colonies from the heteroresistant isolates demonstrated a general shift in the MIC distributions that correlated with the parent isolate AUC measurements (**Figure 3**). Isolate C1814 (AUC 125.8) demonstrated growth of colonies at both 4 µg/mL and 16 µg/mL and had a statistically significant increase in MIC after cefiderocol exposure compared to both susceptible controls PAO1 and C1273 (P-value= 0.024 and 0.017, respectively). Isolates C1526 (AUC 109) and C4587 (AUC 99.6) also demonstrated growth of colonies at 4 µg/mL and 16 µg/mL, with a broadened MIC distribution after cefiderocol exposure. A small number of isolates from the post-exposure population displayed MICs in the 2-4 µg/mL range, compared to 0.5 to 1 µg/mL in the parent population, although these changes were not statistically significant. Colonies from PAO1 were recovered from both the 4 µg/mL and 16 µg/mL cefiderocol growth condition, but the overall distribution of MICs between exposed and unexposed populations was similar and no isolates with an MIC of greater than 1 µg/mL were recovered. The susceptible clinical strain C1273 did not demonstrate any breakthrough growth at a cefiderocol exposure of 16 µg/mL, and all recovered isolates at lower concentrations had an MIC of ≤ 0.5 µg/mL.

**Figure 3.**
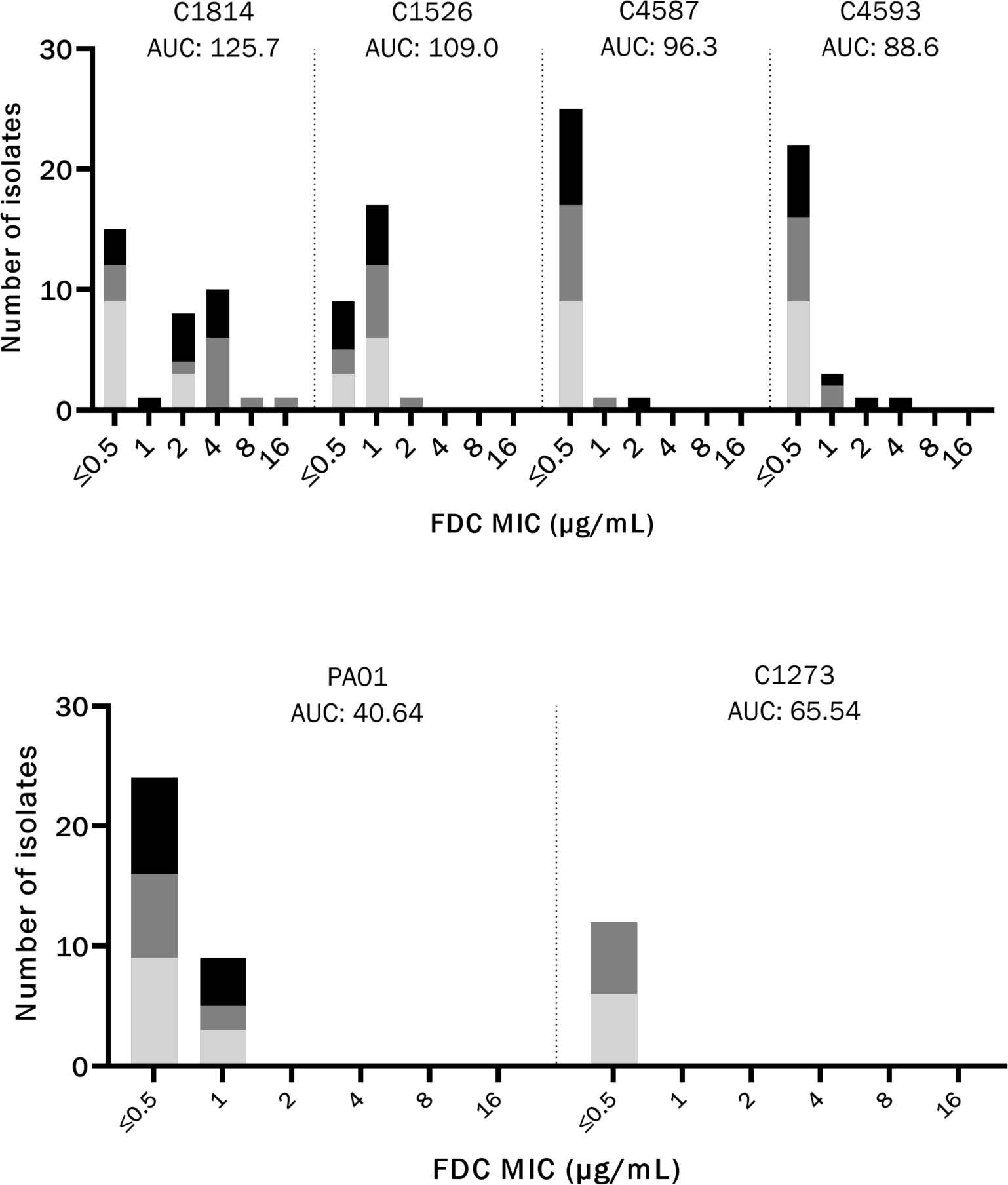
PAP-AUC correlates with a shift in MIC distribution of population after a single FDC exposure. Stacked bar chart with cefeiderocol (FDC) MICs of individual colonies recovered from PAP plates after FDC exposure to 0 µg/mL (bottom, light gray), 4 µg/mL (middle, mid-gray), and 16 µg/mL (top, black). Heteroresistant isolates and AUC are shown in the top panel, with susceptible controls in bottom panel.

### Kirby-Bauer disk diameter correlates with heteroresistance as determined by PAP-AUC

The heteroresistant phenotype is not identified on routine antimicrobial susceptibility testing, and population analysis is a time and labor-intensive method that is not practical for the clinical microbiology laboratory. We noted on disk diffusion testing that heteroresistant isolates were more likely to have breakthrough colonies or dual concentric halos as compared to susceptible isolates, and that these were more pronounced on incubation at 48 hours. Thus, we sought to investigate whether disk diffusion diameters correlated with PAP-AUC results. At 24-hours, the PAP-defined heteroresistant TBDR mutant isolates had significantly smaller KB disk diameters when compared to TBDR mutants with a PAP-defined susceptible phenotype and the wild-type TBDR controls (**Figure 4A**). The mean 24-h KB disk diameter for heteroresistant isolates was 24.5 mm versus an average disk diameter of 29.3 mm for susceptible isolates (P-value = 0.014). The mean 48-h KB disk diameters were similar, with an average of 21.5 mm for heteroresistant isolates versus 27.9 mm for their susceptible counterparts (P-value=0.013, **Figure 4B**). Linear regression was performed by plotting each isolate disk diffusion diameter against the AUCs obtained from the initial PAP screen. Both 24-hour and 48-hour KB disk diameter results correlated strongly with AUC measurements (R^2^= 0.82 and 0.80, respectively), suggesting a simple modification of disk diffusion testing may be a useful screening method to identify heteroresistant isolates.

**Figure 4.**
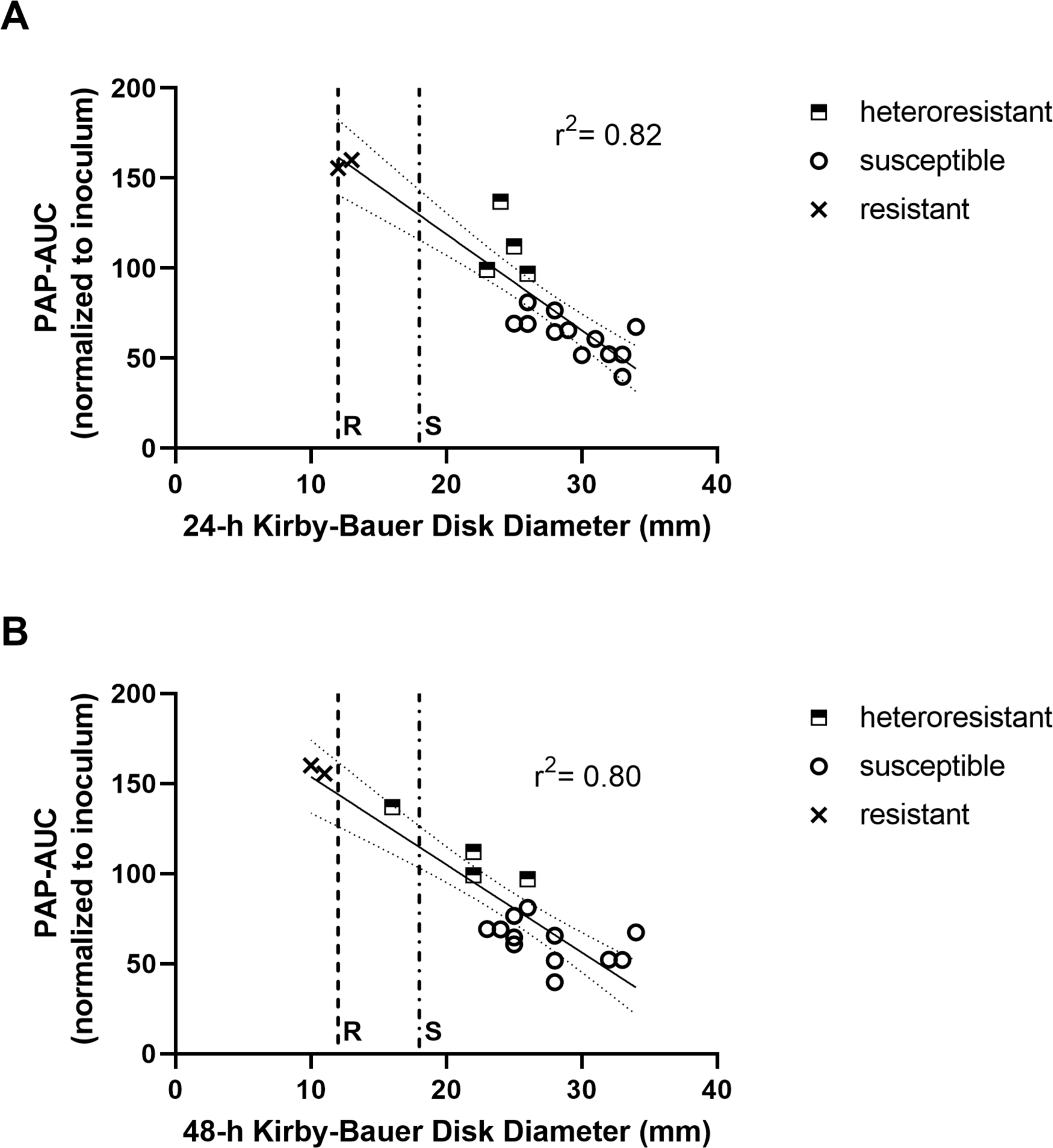
Correlation of heteroresistance phenotype by PAP-AUC to Kirby Bauer disk diameter. AUC values versus Kirby-Bauer disk diameters at **A)** 24-hours and **B)** 48-hours. Resistant (R, dash line) and susceptible (S, dot-dash line) CLSI disk breakpoints indicated. Linear regression with standard error is overlaid on KD disk diameters.

## Discussion

Cefiderocol has *in vitro* activity against many MDR-Gram negative organisms, including *P. aeruginosa*, and has become an increasingly utilized salvage therapy for Gram-negative organisms resistant to most other β-lactam antibiotics (29). Resistance to FDC in *P. aeruginosa*, while uncommon, has been described in association with several factors, including the alteration of the outer membrane TBDRs or the signaling pathways that control their expression, acquisition of exogenous β-lactamases, mutations in penicillin binding protein 3 (PBP3), and changes in the intrinsic PDC enzyme that may alter the hydrolytic activity of the β-lactamase towards FDC (30). We hypothesized that alterations of the TBDR pathway may function similar to other porin mutations, restricting import of the antibiotic to the periplasmic space. While several studies have investigated the presence of TBDR pathway mutations in clinical isolates, these have generally been limited in the number and geographic representation of isolates included (7, 8, 31, 32). In this study we evaluated 498 clinical isolates of *P. aeruginosa* collected across three diverse geographic regions and four unique clinical sites prior to the introduction of cefiderocol. We found pre-existing mutations within the PirA and PiuA/D pathways in up to a quarter of genomes. Importantly, changes likely to have an impact on protein function, including insertion, deletion, or frameshift mutations within the TBDR target genes *pirA, pirS, pirR,* and *piuA/D* were seen at rates up to 5.7% of carbapenem-resistant *P. aeruginosa* isolates from one center, despite no prior cefiderocol exposure. Probable loss of function mutations displayed a geographic predisposition among clinical sites, occurring only in isolates from two different healthcare facilities in Houston, TX. The isolates were from a diverse set of sequence types, suggesting clonal expansion of a single isolate at one center did not appear to explain the geographic distribution of mutations. A number of additional individual SNPs were prominent throughout each center independent of the geographic location of isolates. The specific factors driving selection of mutations in the TBDR genes is unknown. Further studies are needed to elucidate the clinical risk factors and geographic predispositions for isolates with TBDR mutations.

Notably, mutations in any single pathway are rarely linked to overt FDC resistance, suggesting that a combination of factors are needed for the resistant phenotype to emerge. Our data further supports the hypothesis that individual TBDR pathway mutations play a partial or first-step role in the evolution of cefiderocol resistance among clinical isolates. This is consistent with previously published *in vitro* data that suggest the impact of a lone *pirA* mutation on FDC susceptibility is limited, but substantial increases in MIC may occur when changes in PirA occur alongside other TBDR gene mutations (e.g. *piuA/D*) or changes in PDC that may expand its hydrolytic activity against FDC (6, 8, 15). Independently, loss of PiuA/D appears to have a more substantial impact on cefiderocol susceptibility, leading to MIC increases from 0.5 to 8 μg/mL; however, compound mutations involving both the PirA and PiuA cefiderocol TBDR pathways consistently displayed even higher MICs in the ≥16 μg/mL range (15, 16). In this study, the most frequently observed major mutation was a frameshift mutation leading to a premature stop codon in the *pirR* gene, and similar mutations have been described in *P. aeruginosa* of clinical origin (6, 8). Interestingly, expression of *pirA* seems to rely on induction via catechol substrate recognition by PirS, while *piuA* is constitutively expressed in a PirR dependent manner (17).

Thus, loss of a functional PirR would be predicted to impair expression of two of the major TBDRs involved in FDC uptake, and the presence of this mutation in clinical isolates would be of concern for the emergence of resistance. Prior studies have shown that isolates with exposure and resistance to ceftolozane-tazobactam may also display an increased risk for the emergence of cefiderocol resistance, particularly in the setting of both TBDR and *ampC* mutations (7, 8). Consistent with this hypothesis is the fact that the two FDC non-susceptible isolates from this study had *pirR* mutations coupled with either changes in the PDC enzyme (C1418, PDC-205) or an extended spectrum OXA variant (C1408, OXA-15 with G150S substitution) (33). Further investigations are warranted to unravel the prevalence of the combinations of these mutations and their impact on cross-resistance among newer β-lactam/β-lactam inhibitor combinations.

Since there was significant variability in the FDC MICs of isolates with TBDR mutations, we sought to investigate whether changes in TBDRs may be an underlying genetic factor in the development of heteroresistance to FDC. Heteroresistance, or the presence of bacterial subpopulations with reduced susceptibility to FDC, has been described for *P. aeruginosa*, and has been proposed as a potential explanation for higher-than-expected treatment failures against extensively drug resistant isolates(9, 24). In this cohort, heteroresistant isolates were found to be present among strains with TBDR mutations predicted to result in an insertion, deletion, or frameshift mutations, but were not detected in a subset of control isolates with wild-type TBDR genes. Half of the TBDR mutant isolates studied displayed changes in FDC susceptibility: two were non-susceptible to FDC by traditional AST, and 4 of the remaining 10 TBDR mutants displayed a heteroresistant phenotype. The frequency of resistant subpopulations as measured by AUC within heteroresistant isolates directly correlated with *in vitro* emergence of decreased cefiderocol susceptibility after as little as a single drug exposure, emphasizing the potential vulnerability of heteroresistant isolates to adapt upon antibiotic challenge. Further, we provide a comparison of agar dilution methods to detect FDC heteroresistance to a time kill assay in ID-MH broth. Our analyses demonstrate that cefiderocol heteroresistance detected by agar dilution is also captured via growth in iron-depleted media, suggesting that heteroresistance can be reliably assessed via agar dilution methods and is not an artifact of non-physiologic iron content in the growth media. These *in vitro* data emphasize a need for prospective clinical studies that can help define a meaningful AUC cutoff criteria to potentially identify *P. aeruginosa* at high risk for the emergence of cefiderocol resistance on drug exposure.

Finally, our study provides a basis for the further investigation of a clinically feasible method for detecting heteroresistant isolates in the clinical microbiology lab. Importantly, all 4 PAP-classified heteroresistant isolates were classified as susceptible by standard AST methods using CLSI breakpoints, thus current testing methods may not capture the diversity of FDC susceptibility phenotypes in *P. aeruginosa*. Kirby-Bauer disk diffusion studies read at 24 and 48-hours showed a correlation to formal PAP-AUC values. In clinical practice, PAP-AUC is not feasible due to the time and resource requirements of the test. Optimization of techniques such as a heteroresistance cutoff read at 24-hours or 48-hours may be a potential strategy to identify isolates with an increased risk of the emergence of resistance, although further studies are needed using a larger sample size.

In summary, we identified the prevalence of PirA and PiuA/D TBDR pathway mutations across a collection of clinical *P. aeruginosa* isolates with no prior cefiderocol exposure. While TBDR probable loss of function mutations are relatively uncommon, identified in approximately 3% of the overall cohort, we noted a significant variance across geography suggesting that local prevalence may vary substantially (from 0 to 5.7%). Further surveillance is needed as cefiderocol use increases, as this may apply a selective pressure that alters the frequency of these mutations in the population. A second important aspect of our work is that *P. aeruginosa* isolates with disruption of PirR appear to be enriched for a heteroresistant phenotype. These isolates showed a greater degree of survival in the presence of FDC as compared to isolates without changes in the TBDR genes and the laboratory strain PAO1, despite testing susceptible by routine AST methods. Heteroresistant isolates also showed a shift in the MIC distribution of the population after a single FDC exposure, suggesting this phenotype may predispose to an increased risk for the emergence of resistance. Further work is needed to define risk factors for PirA and PiuA/PiuD pathway mutations that appear to arise in select clinical isolates despite no prior cefiderocol exposures and the impact of heteroresistance on clinical treatment outcomes.

## Acknowledgements

This research was supported by the National Institutes of Health, National Institute of Allergy and Infectious Diseases (NIH/NIAID) grant number R21 AI175821 to W.R.M, NIH/NIAID grants K24 AI121296, R01 AI148342, R01 AI134637, and P01 AI152999 to C.A.A., and a training fellowship from the Gulf Coast Consortia, Texas Medical Center Training Program in Antimicrobial Resistance (TPAMR) (NIAID) T32 AI141349 to S.L.E. J.M.M. received funding from the Asociación Nacional de Investigación y Desarrollo (ANID), Fondo Nacional de Desarrollo Científico y Tecnológico FONDECYT regular Grant 1211947.

## Transparency declarations

WRM has received grant support from Merck and royalties from UpToDate. CAA has received royalties from UpToDate. All other authors have no conflicts to disclose.

## Data Availability Statement

Datasets for the whole genome sequencing are available at the National Center for Biotechnology Information (NCBI) under Bioproject Accession numbers PRJNA896240, PRJNA1063198, PRJNA1065430, PRJNA768616.

